# Interpreting single-cell and spatial omics data using deep networks training dynamics

**DOI:** 10.1101/2024.04.06.588373

**Authors:** Jonathan Karin, Reshef Mintz, Barak Raveh, Mor Nitzan

## Abstract

Single-cell and spatial genomics datasets can be organized and interpreted by annotating single cells to distinct types, states, locations, or phenotypes. However, cell annotations are inherently ambiguous, as discrete labels with subjective interpretations are assigned to heterogeneous cell populations based on noisy, sparse, and high-dimensional data. Here, we show that incongruencies between cells and their input annotations can be identified by analyzing a rich but overlooked source of information: the difficulty of training a deep neural network to assign each cell to its input annotation, or annotation trainability. Furthermore, we demonstrate that annotation trainability encodes meaningful biological signals. Based on this observation, we introduce the concept of signal-aware graph embedding, which facilitates downstream analysis of diverse biological signals in single-cell and spatial omics data, such as the identification of cellular communities corresponding to a target signal. We developed Annotatability, a publicly-available implementation of annotation-trainability analysis. We address key challenges in the interpretation of genomic data, demonstrated over seven single-cell RNA-sequencing and spatial omics datasets, including auditing and rectifying erroneous cell annotations, identifying intermediate cell states, delineating complex temporal trajectories along development, characterizing cell diversity in diseased tissue, identifying disease-related genes, assessing treatment effectiveness, and identifying rare healthy-like cell populations. These results underscore the broad applicability of annotation-trainability analysis via Annotatability for unraveling cellular diversity and interpreting collective cell behaviors in health and disease.

## 1 Introduction

Single-cell genomics provides a detailed view of gene expression patterns in individual cells. This level of granularity is used to decode complex biological processes and disease states. It facilitates the identification of distinct types and states of cells[1], the inference of cellular interactions within a tissue or organism[2], and the characterization of temporal dynamics during processes such as development[3], the progression of a disease[4], or treatment-induced response[5].

To organize the vast, noisy, and high-dimensional raw data from single-cell genomics experiments into interpretable patterns, individual cells can be labeled or annotated in terms of their type, state, location, or phenotypic association. The annotation of single-cell genomics data serves as a linchpin in unraveling the intricate details of cellular identities, function, and responses to various stimuli. For example, annotation of cells to cell types is crucial for attributing observed variations in molecular profiles to specific cell types and their function; similarly, labeling of cells as originating from healthy, diseased or treated tissues can aid in discerning subtle changes along disease progression and the effects of therapeutic interventions, thus refining our understanding of disease mechanisms and treatment impacts.

Cell annotations can be either assigned directly based on experimental conditions (e.g. samples originating from patients vs. healthy controls), or inferred by computational analysis of cellular gene expression and additional measured attributes[6]. Annotations of cells e.g. to cell types and states can be inferred based on established marker genes[7, 8], clustering of gene expression profiles[9–11], combinations thereof[12], or in a supervised manner using complementary pre-annotated datasets[1]. Annotations related to localization across spatial or temporal processes can be reconstructed based on prior knowledge of the topology of a biological signal[13, 14], principles of tissue organization[15, 16], trajectory reconstruction[17, 18], or annotation transfer between datasets[19–21].

The computational annotation process, however, can be inherently ambiguous. It is compounded by the need to assign discrete labels, potentially with subjective interpretations, to heterogeneous cell populations based on noisy, sparse, and high-dimensional data. For example, computationally inferred annotations may not capture intermediate states along e.g. immune activation, spatial patterning, or development; similarly, annotations based on experimental conditions may not capture inherent heterogeneity within a cellular population where e.g. a subpopulation of cells in a diseased sample may in fact resemble characteristics of the healthy population. As a result, annotations must often be refined manually based on prior information relevant to a particular system, followed by arduous verification based on independent statistical analyses, expert knowledge, in vitro functional assays, imaging, single-cell qPCR, or mutation data[6]. Even then, due to their inherently ambiguous nature, a substantial subset of the assigned annotations may be fully or partially incongruent with the cells they aim to describe.

Here, we show that the level of congruence between cells and their original annotations carries critical information for downstream interpretation of single-cell genomic data. This information can be used to identify and re-annotate mislabeled cells, identify transient or intermediate cell states, and identify and characterize unique cellular subpopulations such as healthy-like cells within a diseased sample. To characterize such congruence within a cell population relative to a given set of annotations, we exploit an information source that is usually discarded during the training of deep neural networks (DNNs) on such data, namely, the dynamics of the training process itself. Specifically, we do not use DNNs for direct predictions of cell types, states or other types of annotations, as has been done in many recent works[22, 23]. Instead, we focus on the time it takes and the stability with which a DNN learns to predict the original annotation for each cell in the data.

DNNs were shown, in many cases, to have the capacity to memorize practically any training dataset, including noisy data points and inaccurate labels[24]. Although DNNs will eventually learn all input labels or annotations provided to them, empirical evidence suggests a progressive learning pattern across epochs; DNNs tend to first learn data points that were correctly annotated and associated with low noise, and then progress to those characterized by high noise and/or incorrect annotations[25]. Indeed, in the context of image classification, the learning time of an input data point has been successfully used to determine its correct labeling[26, 27]. In the context of NLP datasets, “Data Maps” were introduced to analyze a model’s behavior during training, providing two measures for each data point: the model’s confidence in its input label, and the variability of this confidence across training epochs. This approach has demonstrated that these measures are useful for predicting the reliability of the input label for each data point by its training dynamics, with easy-to-learn data points corresponding to correct input labels, hard-to-learn data points corresponding to erroneous input labels, and moderately-challenging (ambiguous) data points corresponding to ambiguous input labels[28].

We present Annotatability, a method to identify meaningful patterns in single-cell genomics data by quantifying the congruence between a cell and its input annotation, or simply, annotation-trainability analysis. It does so by monitoring the process of training a DNN to predict the input annotations for each cell; and inspecting the confidence and variability with which they are learned (Figure 1a-b). Building on this basic approach, we extend Annotatability with several modules that address higher-level challenges of single-cell genomics analysis, including auditing and rectifying erroneous annotations (Figure 1c-d), identifying ambiguous or intermediate cell states (Figure 1c), identifying cellular communities with shared gene expression profiles and annotation-trainability via graph embedding (Figure 1e-f), and detecting annotation-associated genes (Figure 1g). We demonstrate the utility of Annotatability in different real-world scenarios, including the identification of false annotations of cell types and intermediate cell types in human peripheral blood mononuclear cells (Figure 2); inference and correction of false cell type annotations in spatial transcriptomics of MERFISH mouse hypothalamic preoptic region (Figure 3); inference and embedding of epithelial and mesenchymal cells according to epithelial-to-mesenchymal transition dynamical process (Figure 4); and finally, inference and embedding of pancreatic *β*-cells from diabetic models according to disease-related cell states, with applications for screening for disease-associated genes, evaluation of treatment effectiveness, and sensitive detection of rare subpopulations of healthy-like cells (Figure 5).

**Figure 1.**
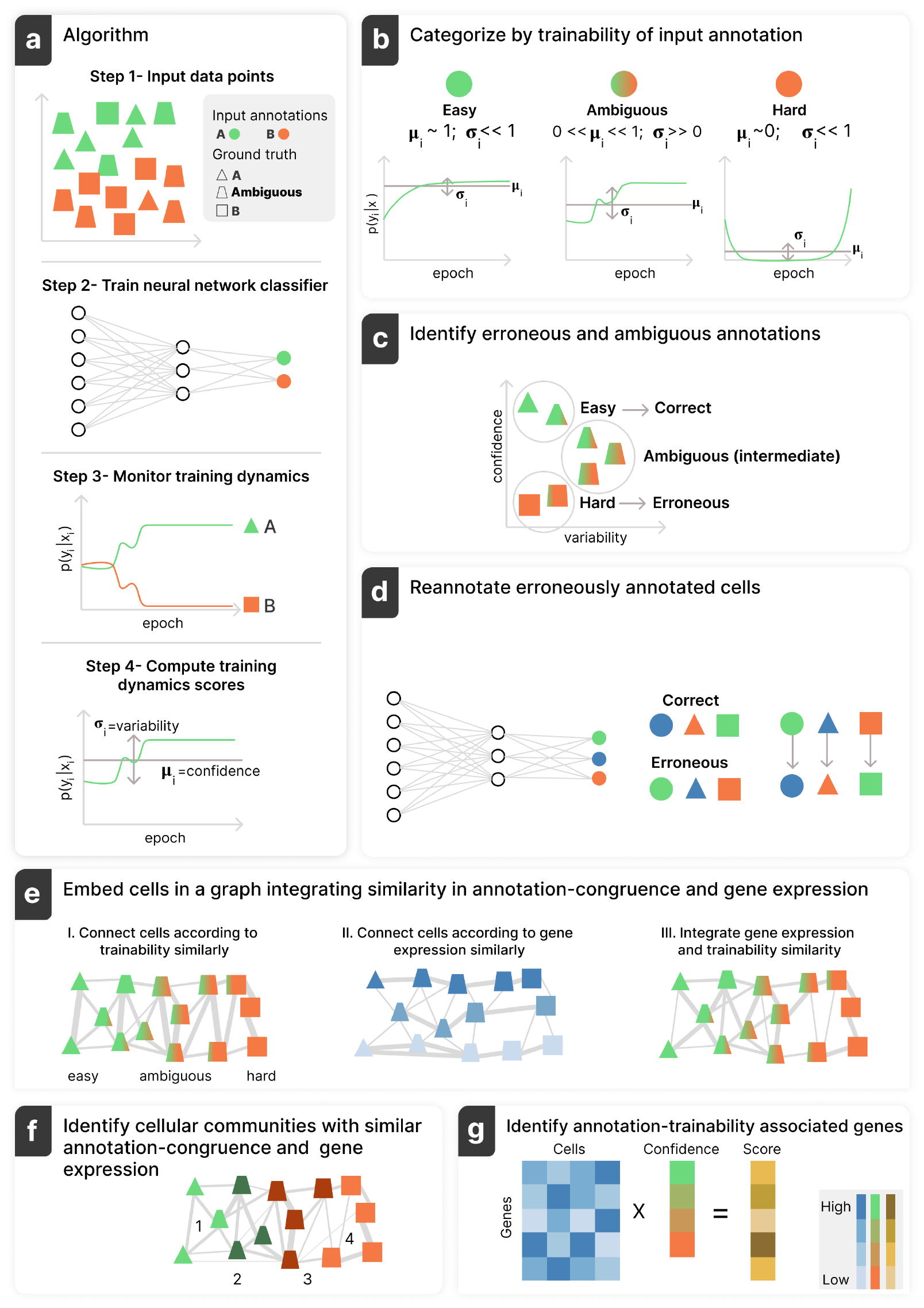
Annotatability schematic workflow. (a) Given an annotated dataset consisting of observations (e.g. single-cell gene expression profiles) and corresponding annotations per cell (e.g. cell types) (Step 1), Annotatability trains a DNN to predict the input annotations (Step 2), analyzes its predictions along the training procedure (Step 3), and computes confidence and variability scores per cell (Step 4). (b, c) Each cell is subsequently classified as easy-to-learn (high confidence/low variability), hard-to-learn (low confidence/low variability), or ambiguous (mid confidence/high variability)(b), corresponding to correctly annotated, erroneously annotated, or ambiguously annotated cells, respectively (c). (d) Annotatability can potentially amend the annotations of cells identified as erroneously annotated using its re-annotation module. (e, f) Annotatability includes a trainability-aware graph-embedding module, which incorporates similarity between cells in both training dynamics statistics as well as gene expression (e), enabling signal-specific downstream analyses (f). (g) Annotatability is also equipped with a training-dynamics-based score that captures either positive or negative association of genes relative to a given annotations, corresponding to a given biological signal.

## 2 Results

### 2.1 Uncovering cell characteristics and tuning annotations in single-cell data using deep learning training dynamics

We developed Annotatability, a framework for annotation-trainability analysis, achieved by monitoring the training dynamics of deep neural networks (DNNs). We use Annotatability to improve the accuracy and resolution of single-cell genomics annotations, identify intermediate cell states, and enable signal-aware downstream analysis. Our working hypothesis, supported by findings in the context of natural language processing and computer vision[26–28], is that along the training process, a DNN first learns decision boundaries between the correctly annotated data points, capturing the general structure underlying the annotated data. As the training process progresses, the network begins to overfit, eventually classifying all data points strictly according to their input annotations, whether they are correct, erroneous, or ambiguous.

Annotatability quantifies these training dynamics as follows. First, it takes an annotated dataset consisting of observations such as gene expression profiles of single cells, and corresponding annotations such as cell types or disease states per cell as input (Figure 1a: Step 1). Second, it trains a DNN, in our case, a simple multilayer perceptron, to predict the input annotation of each cell (Methods; Figure 1a: Step 2). Third, it analyzes the DNN’s predictions along the training procedure (Figure 1a: Step 3). Finally, it computes two scores for each cell[28]: (1) the cell’s confidence score is the mean probability assigned to its input annotation across epochs, and (2) the cell’s variability score is the standard deviation of that probability across epochs (Methods 4.1; Figure 1a: Step 4).

Following the above working hypothesis, we expect learning to become easier, quicker, and less volatile as the fit between a cell and its input annotation increases, and vice versa. Consequently, we expect correctly annotated cells to have high confidence and low variability scores, erroneously annotated cells to have low confidence and low variability scores, and ambiguously annotated cells to have mid-confidence and high variability scores. By setting appropriate thresholds (Methods 4.4), we classify each cell to one of these three categories: correctly annotated (easy-to-learn), erroneously annotated (hard-to-learn), or ambiguously annotated (Figure 1b,c).

Annotatability uses these trainability-based cell classifications for downstream biological analysis in several ways. First, it allows us to focus any downstream analysis solely on correctly annotated cells and study cells that were classified as ambiguously annotated as candidates representing intermediate cell states. It also allows us to identify distinct subpopulations of cells that do not fit their input annotations, such as in the case of healthy-like cells in a diseased sample. Second, the identification of correctly and erroneously annotated cells can be used to amend erroneous annotations; we equipped Annotatability with an optional re-annotation module (Figure 1d), which corrects erroneous annotations by training a DNN solely on correctly annotated cells. The network is then used to re-annotate the entire dataset (Methods 4.5). Re-annotation is particularly useful in contexts where the exclusion of a subset of (incorrectly annotated or ambiguous) cells can pose challenges for downstream analysis, such as in the context of spatial transcriptomic datasets, where such exclusion may lead to spatial gaps and hinder the analysis of spatial configuration and collective cell behavior. Third, Annotatability incorporates a trainability-aware graph-embedding module, utilizing the trainability of a given annotation as a proxy for biological variation, or signal, encoded by the cell. Recent years have seen the development of many scRNA-seq graph-based analysis pipelines, where nodes represent cells and edges between cell pairs are weighted by their proximity in gene expression space[10, 15, 29–31]. Annotatability extends this approach by also considering the similarity of cells’ Annotatability confidence scores when weighting edges (Methods 4.6). This reweighting enhances signals of interest, such as a cell’s developmental stage or disease state, enabling signal-specific downstream analyses (Methods 4.6; Figure 1e-f). Finally, Annotatability is equipped with a training-dynamics-based score that captures either positive or negative association of genes relative to a given biological signal, revealed by their correlation or anti-correlation with the confidence in a particular annotation (Methods 4.3; Figure 1g).

### 2.2 Identifying erroneous annotations and ambiguous cell states in human PBMCs

We begin by demonstrating how annotation-trainability analysis via Annotatability can be used to identify both erroneously and ambiguously annotated cells, the latter potentially corresponding to intermediate cell types. We apply Annotatability to a single-cell RNA-sequencing (scRNA-seq) dataset of human peripheral blood mononuclear cells (PBMCs)[32]. In the original dataset, each cell was pre-annotated to one of eight cell types (Figure 2a). We used Annotatability to train a DNN classifier that predicts cell types from the annotated PBMC scRNA-seq data, and assign confidence and variability scores to each cell (Figure 2b, c).

**Figure 2.**
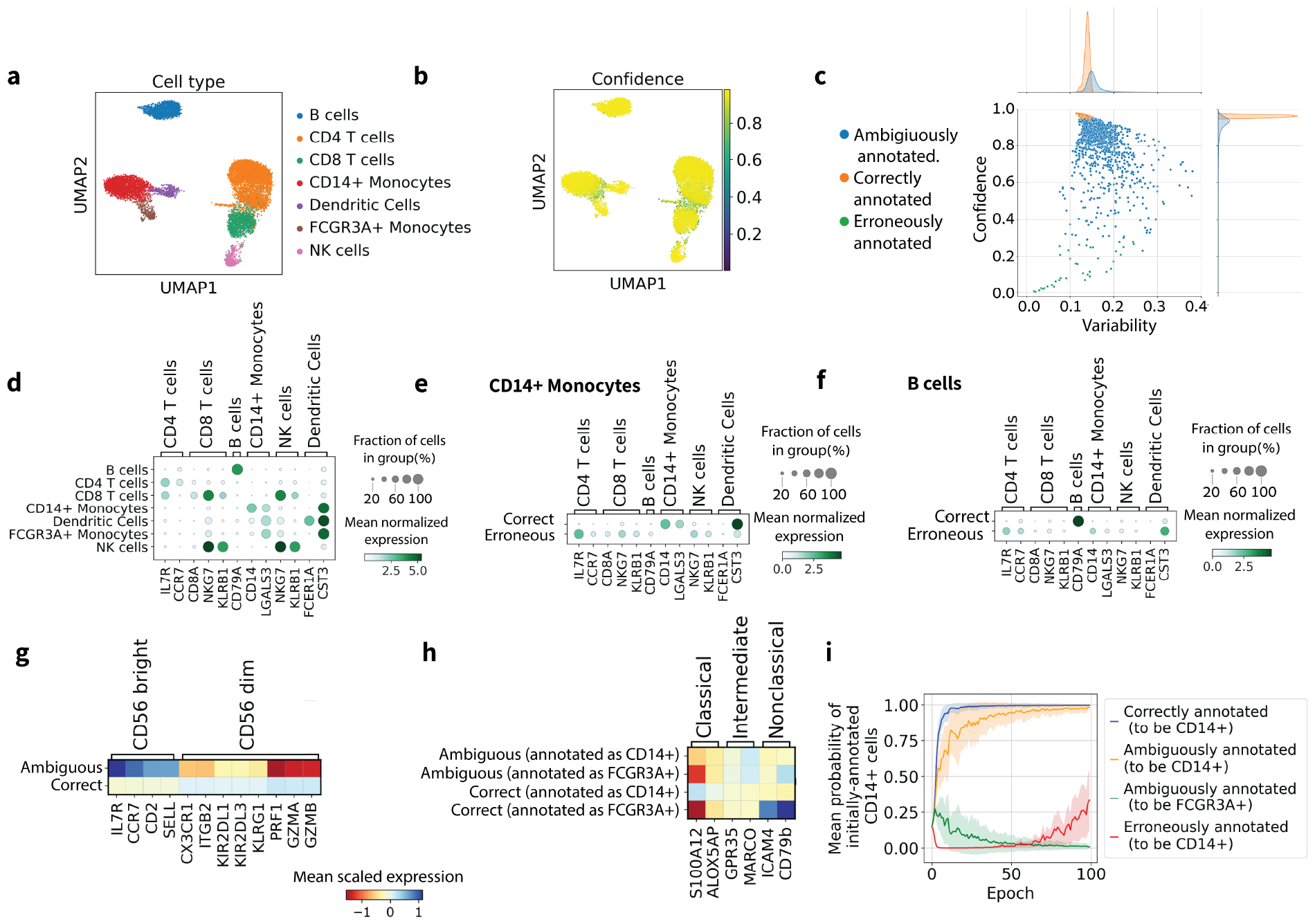
Identification of erroneous and ambiguous annotations in human PBMCs scRNA-seq data[32]. (a, b) 2D UMAPs of cells colored by either the input cell type annotations (a) or the confidence of the cell type annotation inferred by Annotatability. Megakaryocytes were filtered out due to low cell counts (88/11990). (c) 2D confidence-variability map of annotated cells. Subpopulations of cells identified as correctly, ambiguously or erroneously annotated are colored in orange, blue and green, respectively. 1D histograms of each corresponding category appear across the confidence and variability axes. (d) Dot plot of the expression of cell type marker genes in each cluster of the annotated cell types. (e, f) Dot plots of the expression of cell type marker genes of the annotated cell types in either CD14+ Monocytes (e, n=2227) or B cells (f, n=1621) identified as correctly annotated (top row) or erroneously annotated (bottom row). (g) A heatmap of the expression of marker genes of natural killers cells (n=457) subtypes (*CD*56^*bright*^ and *CD*56^*dim*^) identified as ambiguously annotated (top row) or correctly annotated (bottom row). (h) A heatmap of the mean expression of marker genes of monocytes cells (n=2578) subtypes (Classical, Intermediate, and Non-classical) corresponding to cell groups with different initial annotations and trainability characterization. (i) A plot of the mean inferred probability of initially-annotated CD14+ monocytes as a function of training epoch for cells identified as correctly annotated (blue), ambiguously annotated (orange), and erroneously annotated (red) CD14+ cells and ambiguously annotated to be FCGR3A+ cells (green).

We first assess the set of cells classified as erroneously annotated by Annotatability based on their low confidence and low variability scores (Methods 4.4). These erroneously annotated cells indeed exhibit underexpression (or no expression) of marker genes corresponding to their input cell type annotation (Figure 2d; Supplementary Figure 1a, b). For example, cells classified as erroneously annotated CD14+ monocytes underexpress *CD14* yet ∼80% of them express the T cell marker *IL7R* and ∼60% express the NK/T cell marker *NKG7*. Similarly, cells classified as erroneously annotated B cells underexpress the B cell marker *CD79A*, yet a substantial fraction expresses various T cell or dendritic cell markers (Figure 2d-f). Thus, the classification of low-confidence and low-variability cells as erroneously annotated is independently validated by the underexpression of marker genes corresponding to the input cell type annotation, or the expression of marker genes corresponding to other cell type annotations. Additional examples are provided in Supplementary Figure 1).

We followed by assessing whether cells classified as ambiguously annotated indeed correspond to intermediate or otherwise ambiguous cell types. Peripheral blood NK cells are divided into two main subtypes: *CD*56^*dim*^ and *CD*56^*bright*^, comprising the majority and minority of peripheral NK cells, respectively[33, 34]. We found that NK cells classified by Annotatability as correctly annotated express, on average, eight known *CD*56^*dim*^ marker genes[34]. Conversely, NK cells classified as ambiguously annotated express, on average, all four known *CD*56^*bright*^ marker genes[34] that were not filtered out during preprocessing (HVG selection, Methods 4.7; Figure 2g). This classification can be rationalized as follows. First, *CD*56^*bright*^ cells belong to an earlier and less differentiated developmental stage than *CD*56^*dim*^ cells[34]. Second, the gene expression profiles of *CD*56^*bright*^ partially resemble those of T cells (Supplementary figure 1c), and specifically *CD*4+ naive T cells, which share some *CD*56^*bright*^ marker genes (*IL7R, CCR7*[35]; Supplementary Figure 1a); this similarity may hinder the neural network’s ability to classify *CD*56^*bright*^ cells as NK cells.

We also assessed cells classified as ambiguously annotated in the monocytes population. Monocytes are broadly divided into three major subtypes: classical (*CD*14^++^*CD*16^−^), intermediate (*CD*14^++^*CD*16^+^), and nonclassical (*CD*14^+^*CD*16^++^)[36]. However, in the original scRNA-seq data, monocytes were annotated as either classical (*CD*14+) or nonclassical (FCGR3A+), where FCGR3A is a common alias for *CD*16. Thus, we expect intermediate cells in the PBMC dataset to be classified as ambiguously annotated. To support our approach, we used previously established marker genes[32], specifically, the top two highly expressed marker genes for each monocyte subtype that were not filtered out during preprocessing. Indeed, *CD*14+ monocytes classified by Annotatability as ambiguously annotated express lower levels of classical monocyte markers and higher levels of intermediate monocyte markers than *CD*14+ monocytes classified as correctly annotated (classical monocyte markers: mean expression of *S100A12* /*ALOX5AP* is 2.463/0.298 in ambiguous compared to 3.850/0.685 in correctly annotated, p-value of 3.087e-83/1.23e-20 in Wilcoxon test; intermediate monocyte markers: mean expression of *GPR35* /*MARCO* is 0.149/0.413 in ambiguous compared to 0.105/0.202 in correctly annotated, p-value of 0.069/5.34e-08 in Wilcoxon test) (Figure 2d, h). Similary, FCGR3A+ monocytes classified as ambiguously annotated express lower levels of non-classical monocyte markers and higher levels of intermediate monocyte markers than FCGR3A+ monocytes classified as correctly annotated (non-classical monocyte markers: mean expression of *ICAM4* /*CD79b* is 0.118/0.926 in ambiguous compared to 0.655/1.620 in correctly annotated, a p-value of 2.23e-11/1.40e-06 in Wilcoxon test; intermediate monocyte markers: mean expression of *GPR35* /*MARCO* is 0.168/0.502 in ambiguous compared to 0.083/0.167 in correctly annotated, p-value of 0.173/0.002 in Wilcoxon test) (Figure 2d, h).

Having validated Annotatability’s categorization of cells to either correctly, erroneously or ambiguously annotated cells, we next directly evaluate differences in training dynamics among these three categories as a function of the learning epoch. We focus on the group of cells originally annotated as classical (*CD*14+) monocytes. For each epoch, we compute the mean probability that cells in each category are recognized as classical monocytes by the neural network. We identify three distinct regimes along training: early (0-5 epochs), middle (5-50 epochs), and late (50-100 epochs). For cells in the correctly annotated category, the mean inferred probability to keep the classical (*CD*14+) annotation increases to a value close to 1.0 already during the early epoch regime, and remains high throughout the training process (Figure 2i, blue). In that sense, correctly annotated cells are easy to learn. For cells in the incorrectly annotated category, the mean inferred probability decreases to a value close to 0.0 during the early regime, remains low throughout the middle regime, yet gradually increases during the late regime, reaching a value of 0.25 (Figure 2i, red). In that sense, erroneously annotated cells are hard to learn, but the network begins to ‘memorize’ their incorrect annotations during the late regime, which is therefore also termed the overfitting regime. For cells in the ambiguously annotated category, the mean inferred probability to keep the classical (*CD*14+) annotation increases only gradually during the early and middle regimes, saturating only upon reaching the late regime (Figure 2i, orange). For the same set of cells, the mean inferred probability that they are in fact non-classical monocytes (FCGR3A+) increases during the early regime, but reverts to 0.0 during the middle regime. This example demonstrates how the training dynamics patterns of ambiguously annotated cells differ from those of correctly and erroneously annotated cells, and why they can inform an intermediate transcriptional state that is neither classical nor non-classical (Figure 2i).

### 2.3 Re-annotating spatial transcriptomics data

After identifying erroneously annotated cells using Annotatability, one option is to filter them from their corresponding single-cell genomics dataset, based on the assumption that the remaining cells retain sufficiently rich information. However, in some cases, discarding all misclassified cells may disrupt the biological interpretation of the data. For example, in spatially-aware single-cell transcriptomic measurements[37], preserving complete, gap-free spatial maps of the measured regions is crucial for correctly interpreting spatial gene expression and for inferring associated collective cell behavior. Furthermore, the annotation of spatial data poses unique challenges compared with non-spatial data, as many of the spatially-aware transcriptomics experimental methods are limited in the number of genes that are measured, and/or in their spatial resolution (e.g. each spatial location represents averaged measurements of multiple cells) [38].

We therefore equipped Annotatability with a re-annotation module for the robust re-annotation of cells identified as erroneously annotated (Methods 4.5). We evaluated the capability of Annotatability to correctly re-annotate spatial transcriptomics data for a MERFISH dataset of mouse hypothalamic preoptic region[39] (Figure 3a,b). We first employed Annotatability to identify erroneously annotated cells, monitoring the learning dynamics of a DNN trained on the original input annotations, and computing confidence and variability scores for each cell (Figure 3c,d).

**Figure 3.**
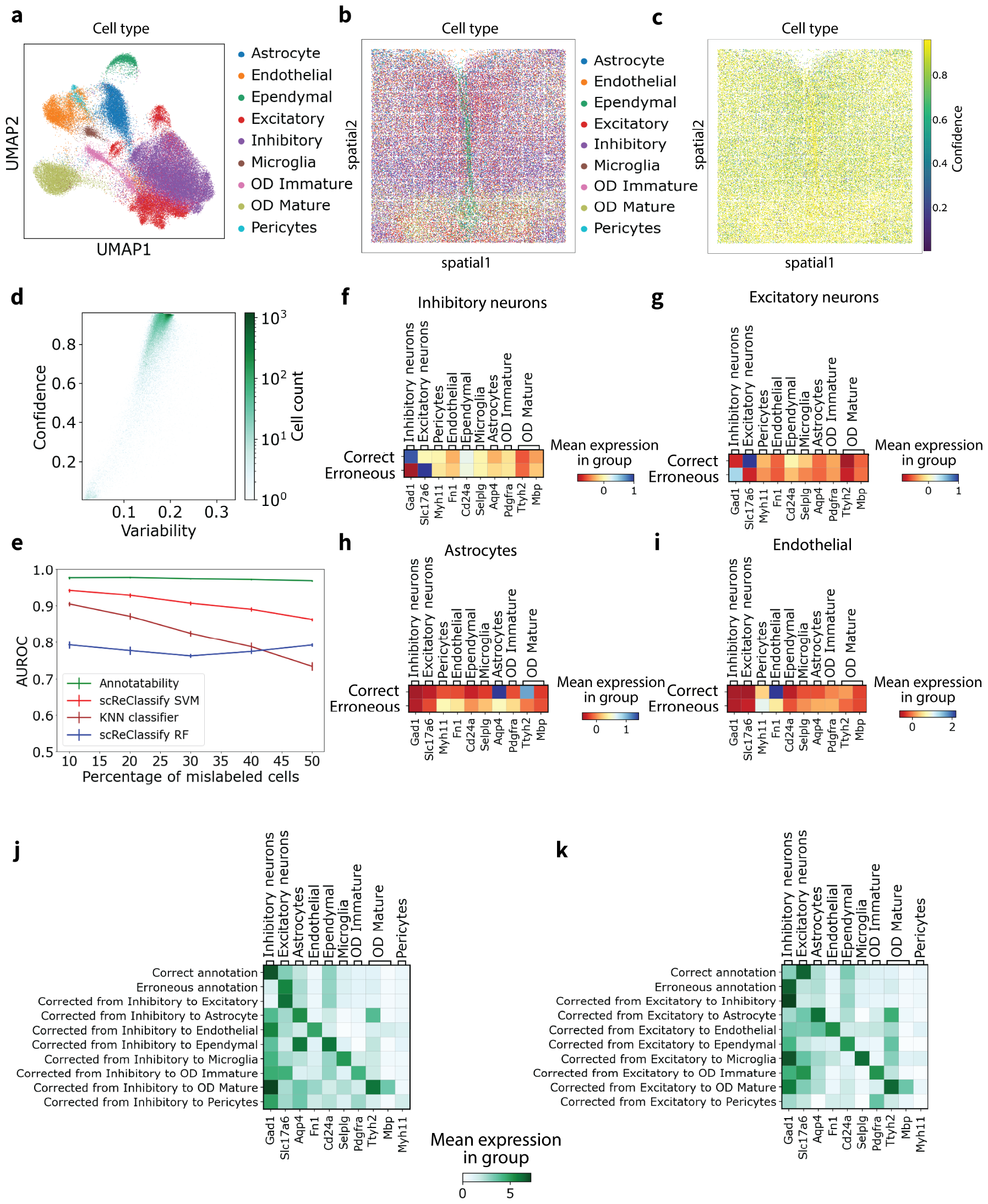
Re-annotation of erroneous annotations in spatial transcriptomics MERFISH dataset of mouse hypothalamic preoptic region[39] (n=73642). (a) 2D UMAP of cells colored by cell type annotations. (b, c) Spatial scatter plot of MERFISH data colored by either cell type annotations (b) or cell type annotation confidence inferred by Annotatability (c). (d) 2D confidence-variability map of the annotated cells. (e) AUCROC vs percentage of mislabeled cells, for four different methods: Annotatability(green), scReClassify with SVM classifier (red), KNN classifier (brown) and scReClassify with random forest classifier (blue). (f-i) Heatmaps of the mean expression of marker genes of the annotated cell types, in Inhibitory neurons (f, n=24761), Excitatory neurons (g,n=11757), Astrocytes (h, n=8393), or Endothelial cells (i, n=5749) identified as correctly annotated (top row) or erroneously annotated (bottom row). The cell type marker gene[39]: *Gad1* (Inhibitory neurons), *Slc17a6* (Excitatory neurons), *Myh11* (Pericytes), *Fn1* (Endothelial), *Cd24a*(Ependymal), *Selplg* (Microglia), *Aqp4* (Astrocytes), *Pdgfra* (OD Immature), *Ttyh2, MBbp* (OD Mature). (j, k) Heatmaps of the mean expression of marker genes of cells that were initially annotated as either Inhibitory neurons (j) or Excitatory neurons (k), and identified by Annotatability as either correctly annotated (first row) or erroneously annotated (second row), as well as their expression after re-annotation by Annotatability (next rows).

To benchmark our approach, we first used a semi-synthetic dataset, created by randomly perturbing the annotation of varying fractions of cells (10%-50%), with the underlying assumption that the majority of cells were correctly classified in the original data. We then used Annotatability to identify the erroneously annotated cells in the perturbed data. We compared the performance of Annotatability with k-nearest neighbors classifier, as well as scReClassify, a state-of-the-art method for the identification of misclassified cells[40] based on either a random forest (RF) or a support vector machine (SVM) classifier. Annotatability consistently outperformed all three baselines in distinguishing between correctly and erroneously annotated cells, its advantage increasing with the fraction of synthetically-misclassified cells (AUROC for 10%/50% misclassified cells: 0.977/0.969, 0.947/0.860, 0.797/0.787, 0.911/0.727 for Annotatability, scReClassify SVM, scReClassify RF, and KNN, respectively; Figure 3e).

Similarly, Annotatability outperforms baselines in identifying misclassified cells in the context of three additional spatial datasets, including the 10x Genomics Visium dataset of Coronal section of mouse brain[41], the 4i dataset of sub-cellular, human tissue cell culture[42], and a seqFISH mouse embryonic dataset[43] (Supplementary Figure 2). The relative advantage of Annotatability with respect to baselines in identifying erroneous annotations is further increased in datasets with elevated noise levels (Supplementary Figure 2).

Progressing to a nonsynthetic setting, we inspected cells identified as erroneously annotated in the unperturbed hypothalamic MERFISH dataset. Of the 62,880 cells in the dataset, 1,138 were flagged by Annotatability as erroneously annotated (Figure 3f-i;, Supplementary Figure 2a-d) of which the majority were initially annotated as either inhibitory neurons (587/23687, Figure 3f, j) or excitatory neurons (314/11366, Figure 3g, k). To re-annotate these erroneously annotated cells, we next trained Annotatability’s re-annotation module (Methods 4.5). We then used the trained re-annotation classifier to predict annotations for the set of originally erroneously annotated cells (Methods 4.5). Indeed, following this procedure, the annotations of cells identified as erroneously annotated were transformed into annotations consistent with their marker gene characterization, based on established marker genes for the different cell types[39] (Figure 3j, k, Supplementary Table 1). For example, the mean normalized expression of *Gad1*, a marker gene for Inhibitory neurons[39], is 6.830 in the correctly annotated Inhibitory neurons, 1.458 in the erroneously annotated Inhibitory neurons, and 6.795 in cells re-annotated as Inhibitory neurons. Similarly, the mean expression of *Slc17a6*, a marker gene for Excitatory neurons[39], is 6.130 vs. 1.961 in cells identified as correctly vs. erroneously annotated Excitatory neurons, respectively. Following re-annotation, *Slc17a6* mean expression recovered to 6.165 for cells re-annotated as Excitatory neurons. More generally, for cells that Annotatability identifies as erroneously annotated, marker gene expression, on average, aligns with the new cell type annotations following re-annotation, rather than the original input annotations. (Supplementary Table 1).

Thus, error detection based on training dynamics combined with retraining on a subset of highconfidence/low-variability cells enables robust re-annotation of spatial transcriptomics data.

### 2.4 Uncovering the epithelial to mesenchymal pseudo time trajectory

Going beyond reasoning about annotations of static cell properties and their interpretation, we next demonstrate that annotation-trainability analysis can unveil dynamic trajectories and temporal shifts in cellular states. We illustrate this concept using single-cell data related to the epithelial-to-mesenchymal transition (EMT). EMT is the process through which epithelial cells convert to a mesenchymal state, marked by the loss of cell–cell junctions and cell polarity. The EMT occurs during several biological processes such as embryonic development, regeneration, and wound healing[44]. It also plays a role in abnormal or disease processes, including organ fibrosis, tumor progression with metastatic expansion, and the generation of tumor cells with stem cell properties; the latter plays a major role in resistance to cancer treatment[45].

Traditionally perceived as a binary transition between the E (epithelial) and M (mesenchymal) states[46], the EMT has been conceptually generalized to a continuous temporal transition through intermediate cellular phenotypes, observed along the EMT in the contexts of development and cancer[46–48]. Computationally, these intermediate states can be inferred using techniques such as graph-based clustering and pseudotime reconstruction[46]. However, graph-based clustering that relies solely on gene expression, as commonly done[10], can be sensitive to sources of variation other than the EMT signal, such as treatment effects, as we show below. In contrast, we will demonstrate that Annotatability allows us to construct a trainability-aware gene expression graph that specifically enhances the EMT signal encoded in the data. This enhancement allows for a more in-depth exploration of gene expression patterns and underlying processes driving the EMT.

We used Annotatability to analyze scRNA-seq datasets of MCF10A and HuMEC cell lines[49], which serve as models for benign mammary epithelial cells. Our analysis included both untreated cells (mock condition) and cells treated with the key regulatory cytokine TGF-*β*, which induces the EMT process[49] (Figure 4a), among other effects[50]. To generate initial input labels for Annotatability, we clustered the cells based on their gene expression using Louvain clustering and pre-annotated the cells in each of the resulting 22 clusters based on known marker genes associated with either the epithelial state (*CDH1, CRB3, DSP* [49]) or the mesenchymal state (*VIM, FN1, CDH2* [49]) (Figure 4b). Following standard preprocessing (Supplementary 4.7) and without requiring batch integration, we then applied Annotatability by training a DNN classifer to predict whether a cell is in an epithelial or mesenchymal state, followed by computing the corresponding training dynamics scores (Methods 4.1). We observed a wide spread in the distribution of both the confidence and the variability scores for both the mock and the TGF-*β*-treated cells (Figure 4c). We used Annotatability to refine the initial clustering of cells by identifying erroneously annotated cells (supported by the E/M marker genes; Figure 4d), and filtered these from further analysis.

**Figure 4.**
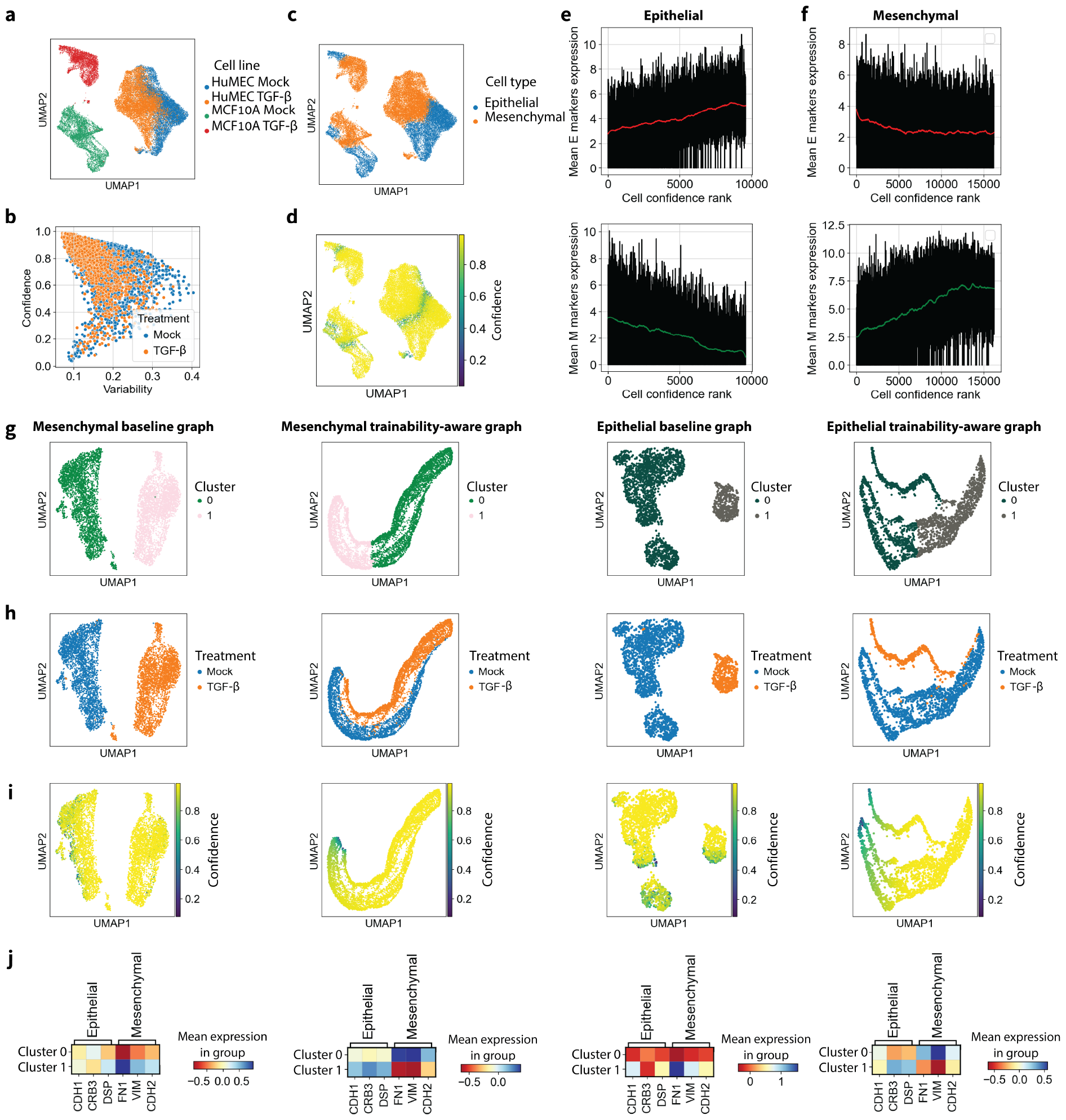
Capturing the epithelial to mesenchymal transition via trainability-aware graph analysis, based on EMT scRNA-seq data[49](n=25806). (a, b) 2D UMAPs of cells colored by either cell line and treatment (a) or cell phenotype. (c) 2D confidence-variability map of the annotated cells, colored by treatment. (d) Heatmaps of the expression of epithelial and mesenchymal marker genes, in initially-annotated epithelial state cells identified as correctly or erroneously annotated (first and second row, respectively), and initially-annotated mesenchymal state cells identified as correctly or erroneously annotated (third and fourth row, respectively). (e, f) Mean expression of epithelial (top row) and mesenchymal (bottom row) marker genes as a function of the inferred confidence rank in epithelial (e) and mesenchymal (f) states. (g-i) 2D UMAPs of (left to right columns): baseline expression graph of Mesenchymal annotated cells (from the MCF10A cell line), trainability-aware graph of Mesenchymal annotated cells, baseline expression graph of Epithelial annotated cells, and trainability-aware graph of Epithelial annotated cells. The graphs are colored by (top to bottom rows): Louvain clustering, treatment, inferred confidence by Annotatability. (j) Heatmaps of scaled and centered expression of Epithelial and Mesenchymal marker genes in cells belonging to the two clusters as in (g). The columns, from left to right, correspond to the graphs as in (g-i).

We hypothesized that the confidence score assigned to each cell by Annotatability reflects the underlying gradual E-M transition and can therefore serve as a proxy for the cell’s state along the EMT process. Indeed, the confidence score of a cell is correlated to its similarity to the two end states (E/M) of the EMT process (Figure 4e, f). Specifically, the mean expression of epithelial marker genes increases (decreases) monotonically with the confidence score of cells originally labeled as epithelial (mesenchymal) (Spearman correlation: 0.414 and −0.453, respectively). Similarly, the mean expression of mesenchymal marker genes increases (decreases) monotonically with the confidence score of cells originally labeled as mesenchymal (epithelial) (Spearman correlation: 0.545 and −0.206, respectively).

To focus on the EMT signal, we applied our trainbability-aware graph embedding algorithm (Methods 4.6) to the MCF10A cell line scRNA-seq data, individually for the Epithelial and Mesenchymal cells, followed by Louvain clustreing of each graph (analogous results were obtained for the HuMEC cell line; Supplementary Figure 3). The trainability-aware graph, integrating information about both training dynamics statistics and gene expression, indeed captures the progression of cells along the EMT dynamical process, as the confidence in the input E/M labels changes gradually along the graphs (Figure 4g-i). This is in contrast to a graph constructed based on gene expression alone, resulting in a graph structure which captures treatment effects (Mock vs. TGF-*β*) instead of the EMT process (Methods 4.6; Figure 4g-i, Supplementary Figure 3a-c). Specifically, in the mesenchymal (epithelial) baseline expression graph, 0.98% (0.99%) of cells in one cluster and 0.99% (0.99%) of cells in the second cluster were treated with Mock or TGF-*β*, respectively. This is in contrast to 0.68% (0.67%), and 0.79% (0.95%) of cells in each of the corresponding clusters in the trainability-aware graph which were treated with Mock or TGF-*β*, respectively. Furthermore, E/M marker genes are differentially expressed at different stages along the trainability-aware expression graph structure (and not along the expression graph structure), further supporting the correspondence between the trainability-aware graph and the EMT process (Methods; Figure 4j, Supplementary Figure 3d).

### 2.5 Inference of disease-related cell states and treatment responses

Having established the utility of annotation-trainability analysis for dynamical processes, we next use it to infer disease states for different cellular communities, identify healthy-like cells in diseased samples, identify disease-associated genes, evaluate treatment effectiveness, and delineate individual cell responses. We analyzed scRNA-seq data collected from pancreatic islets isolated from both healthy mice and from multiple low-dose streptozotocin (mSTZ)-induced diabetic models[51], some of which were treated with various compounds, and some with vehicle.

STZ is a naturally occurring diabetogenic compound that targets and destroys insulin-secreting pancreatic *β* cells, leading to reduced insulin levels and elevated blood glucose levels[52–54]. In the mSTZ-induced diabetic models, STZ is applied in multiple low doses, partially degrading *β*-cell activity[55], where the level of degradation may differ between individual cells[56]. To quantify the damage to individual cells extracted from mSTZ-induced diabetic models, we evaluate their trainability for a disease label against the backdrop of control cells annotated as healthy. We applied Annotatability to the islets dataset across all primary cell types of pancreatic islets (*α, β*, and *δ* cells), and computed corresponding confidence and variability scores (Methods 4.1). To independently assess disease progression, we monitored the expression of nine associated marker genes[57] which were retained during the highly variable gene selection stage of preprocessing 4.7; disease progression is characterized by diminished expression of *β*-cell activity and maturation markers (*Ins1, Ins2, Slc2a2, Trpm5, G6pc2, Slc30a8, Ucn3*) and elevated expression of *β*-cell de-differentiation markers (*Aldh1a3, Serpina7*).

#### Inferring disease states

We first assess whether training dynamics inform the levels of disease progression across all mSTZ cells, without yet considering treatment variations. We hypothesized that among the mSTZ cells, the most severely damaged cells are the easiest to learn, and the least severely damaged cells are the hardest to learn. Indeed, high confidence in the disease annotation is associated with diminished expression of *β*-cell activity markers like *Ins1* (Figure 5a, Supplementary Figure 4a) or elevated expression of *β*-cell de-differentiation markers like *Serpina7* (Supplementary Figure 4b,c), and vice versa. To refine the classification of mSTZ cells by disease level, we constructed a trainability-aware graph (Methods 4.6) followed by Louvain clustering, tuning the clustering resolution to guarantee an output of three clusters (Figure 5b). The three clusters can be distinguished by the expression patterns of the nine disease progression markers (Figure 5b-d; Supplementary Figure 4d-f): a disease cluster expressing low levels of activity and maturation markers and high levels of de-differentiation markers; a healthy-like cluster expressing high levels of activity and maturation markers and low levels of dedifferentiation markers, resembling those in the healthy control; and an intermediate cluster expressing intermediate levels of all markers relative to the disease and healthy-like clusters. We verified that the difference in the expression of disease-associated genes across clusters indeed corresponds to changes in the training dynamics scores; specifically, decreasing confidence in the disease annotation along the disease, intermediate and healthy-like clusters (Figure 5e; Supplementary Figure 4g)). The clustering quality is robust with respect to clustering granularity, consistently manifesting a graded progression from a disease-like to a healthy-like expression pattern of marker genes when the cells are divided to three, four, or five clusters (Figure 5f; Supplementary Figure 4h-m). Thus, the clustering of the trainability-aware graph provides a refined quantification of the damage to individual mSTZ *β*-cells.

**Figure 5.**
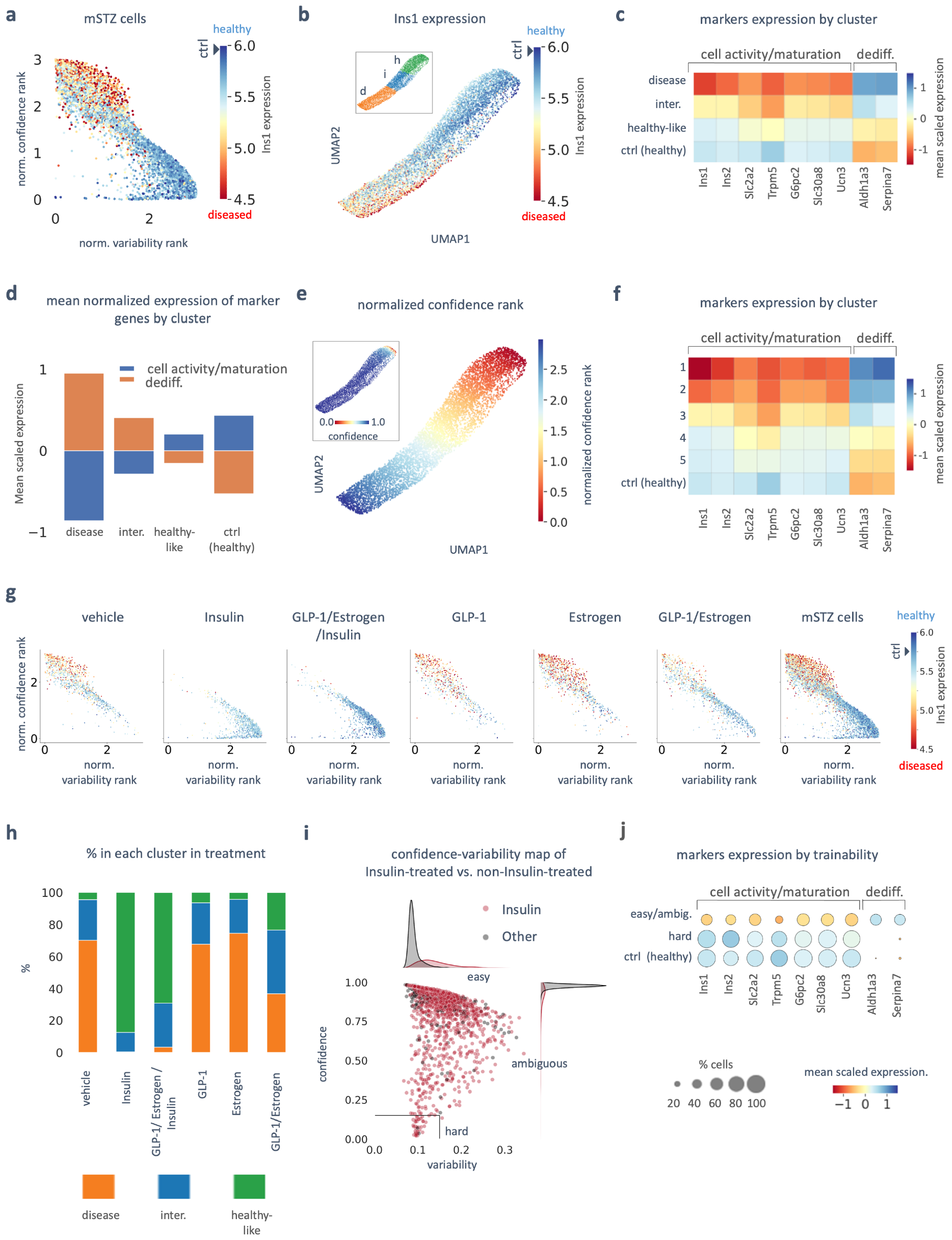
Inferring disease progression, state heterogeneity and treatment responses, based on pancreatic islets scRNA-seq data[51](n=32888). a) 2D map of the normalized ranks of the confidence and variability scores, computed across all mSTZ cells (regardless of follow-up treatment) with respect to a disease annotation, colored by *Ins1* expression. b) 2D UMAP showing the trainability-aware graph colored by *Ins1* expression; inset: the same UMAP colored by the Louvain clustering of the graph; h - healthy-like, i - intermediate, d - disease. c) Heatmap showing mean scaled expression of 9 marker genes for beta cell activity and maturation and beta-cell de-differentiation in the Louvain clusters obtained from the trainability-aware graph, and in the healthy control. d) Bar plot showing mean scaled gene expression for all beta cell activity/maturation markers (blue) and de-differentiation markers (orange) in the Louvain clusters and the healthy control.

#### Identifying disease-associated genes

We next use annotation-trainability analysis to identify genes associated with the disease state of individual mSTZ *β-*cells, in the sense that their expression levels are correlated (positively-associated) or anticorrelated (negatively-associated) with the confidence score assigned to each cell’s disease annotation (Methods 4.3; Figure 1g). Among the five top-ranking genes that are positively-associated with a disease state, we identified known *β*-cells de-differentiation marker genes, including *Gpx3* (Glutathione peroxidase 3)[58], *CD81* [59], *Aldh1a3* [57], and *Aqp4*, which have all been found to be upregulated in *β*-cells from mSTZ-induced diabetic models[59]; and mitochondrial respiratory chain complex I component *Ndufa1*, which is implicated is glucose metabolism regulation[60]. Among the five top-ranking genes that are negatively-associated with a disease state, we identified known *β*-cell activity markers *Trpm5* [57] and *Ins1* [57]; mitochondrial respiratory chain complex IV component *Cox6a2*, which has been found to be downregulated in diabetic islets[61]; *Fam151a*, which is also downregulated in diabetic pancreas[62]; and *Tmem215*, whose function and possible association with diabetes should be further investigated. The full list of disease-association gene scores is provided in Supplementary Table 2.

#### Evaluating treatment effectiveness and individual cell responses

We next apply annotationtrainability analysis to characterize the effectiveness of different diabetes treatments in the mSTZ dataset, which includes six subgroups of mSTZ diabetic models, each treated with a different compound or combinations thereof : vehicle, Insulin, Insulin/GLP-1/Estrogen, GLP-1, Estrogen, and GLP-1/Estrogen. We ranked all mSTZ cells by their confidence and variability scores (Figure 5a), and analyzed the distribution of score ranks in each treatment subgroup (Figure 5g). In principle, the observed heterogeneity in disease progression among mSTZ cells (Figure 5a-f) could reflect treatment-independent heterogeneity, treatment-dependent heterogeneity, or both. To evaluate treatment-independent heterogeneity, we inspected the vehicle-treated subgroup, which serves as a negative control. While most cells in this subgroup were assigned high-ranking confidence scores and low-ranking variability scores, we identified a sizable subgroup with low-ranked confidence scores and high-ranked variability scores (Figure 5g, left panel), indicating lower congruence with a disease annotation. This subgroup is enriched for cells expressing healthy-like levels of *Ins1* (Figure 5a,g, left panel; Supplementary Figure 4a, left panel) and low levels of *Serpina7* (Supplementary Figure 4b-c, left panels). Thus, cellular heterogeneity in disease progression is at least partially independent of treatment, and can be detected using variation in training dynamics scores.

We followed by assessing treatment-dependent heterogeneity, that is, the effectiveness of the different treatments. For the two subgroups that were treated with insulin, either alone or in combination with GLP-1 and Estrogen, the distribution of the confidence and variability scores skewed notably toward lower confidence scores, along with high expression of the *Ins1* gene and low expression of the *Serpina7* gene; in contrast, for the subgroups treated with only GLP-1 or Estrogen, the distribution resembled the vehicle-treated control, and for the subgroup treated with GLP-1/Estrogen, it resembled an intermediate between the insulin-treated and the vehicle-treated cells (Figure 5g; Supplementary Figure 4a-c). Indeed, cells treated with Insulin were distributed nearly exclusively between the intermediate (20.06%) or healthy-like (77.3%) clusters in the trainability-aware graph; in contrast, cells treated with either GLP-1 or Estrogen were overrepresented in the disease cluster (67.79% and 74.55%, respectively), in proportions similar to the vehicle-treated cells (70.17%), with most remaining cells being assigned to the intermediate cluster (GLP-1 25.69%, Estrogen 21.13%, vehicle 25.29%); and cells treated with both GLP-1 and Estrogen were overrepresented in the intermediate state (39.69%) compared with the vehicle, but only a small fraction were assigned to the healthy-like cluster (23.44%) (Figure 5h; Supplementary Figure 4n)).

Upon scrutinizing the raw confidence and variability map, we identified a small subpopulation comprising 43 cells that are particularly hard to learn due to their low variability and confidence levels (confidence<0.15; variability<0.15). Among these, 41 cells were subjected to Insulin or GLP-1/Estrogen/Insulin treatments (Figure 5i). The exceptionally low variability and confidence scores observed in this subpopulation suggest strong incongruence with respect to a disease annotation. Indeed, the cells from this hard-to-learn subpopulation resembled the control healthy group across all nine marker genes (Figure 5j). The preponderance of insulin-treated cells in the healthy-like cluster, together with the nearly complete absence of cells not treated with insulin from the hard-to-learn subset, may imply that insulin treatment is exceptionally effective at restoring a nearly-healthy cell state, which cannot be explained by treatment-independent heterogeneity in mSTZ cells. Moreover, it implies that annotation-trainability analysis is sufficiently sensitive for identifying a rare subpopulation of nearly-healthy cells.

We conclude that treatments of mSTZ mice that include Insulin transform a substantial portion of their *β*-cells into a nearly-healthy state, in contrast with the treatments that do not include insulin. Moreover, annotation-trainability analysis can be used not only for inferring disease state of individual cells, but also for evaluating treatment effectiveness and individual cell responses to the same treatment.

Figure 5. : e) 2D UMAP of the trainability-aware graph colored by the normalized ranks of the confidence score, computed across all mSTZ cells; inset: same UMAP colored by the raw confidence scores. f) Same heatmap as in (c) for five clusters; the number of clusters was increased by tuning the Louvain clustering hyperparameters; see Supplementary Figure 4h-m. g) Same as in (a), split by the different treatments. h) Normalized stacked bar plot showing the share of cells from each Louvain cluster for each treatment. i) 2D confidence-variability map of all mSTZ cells either treated with Insulin (alone or with other compounds; red) or not treated with Insulin (gray). The rectangle indicates the 43 hard-to-learn cells, of which 41 were treated with Insulin (variance <0.15 confidence <0.15); 1D histograms of each corresponding category appear across the confidence and variability axes. j) Dot plots showing the mean scaled expression of the same marker genes as in (c), for either the easy-to-learn + ambiguous mSTZ cells (top row), the hard-to-learn mSTZ cells (middle row), or the control healthy cells (bottom row).

## 3 Discussion

Annotations of single cells to distinct types, states, locations, or phenotypes enrich raw genomic data with descriptive information that provides context, explanation, and detail, thus forming a crucial starting point for its organization and interpretation. However, annotations also force heterogeneous cell populations into rigid molds, whose interpretation is inherently subjective, particularly considering that the annotation relies on noisy, sparse and high-dimensional measurements. In this study, we introduced Annotatability, a method to identify meaningful patterns in single-cell genomics data via annotation-trainability analysis, which taps into the commonly neglected signal encoded in a deep neural network’s training dynamics.

A cell can be incongruent with its input annotation due to several different reasons, from simple assignment errors to fundamental ambiguities associated with underlying biological processes. Based on this observation, Annotatability addresses both routine and fundamental challenges in interpreting single-cell and spatial genomics datasets. First, it can be used for auditing and rectifying annotations in routine single-cell analysis pipelines, by identifying erroneously annotated cells followed by filtering (Figure 2) or re-annotating (Figure 3) these cells based on specific research needs, such as retaining as many cells as possible in analysis of spatial transcriptomics data. Second, it reveals intermediate cell types or states (Figure 2) through the ambiguity in trying to assign them to their input annotations during training. Third, it is used for higher-level downstream analysis tasks, such as delineating temporal cell trajectories (Figure 4), characterizing cellular diversity in diseased tissue, and assessing treatment effectiveness and individual cell responses (Figure 5). For these downstream tasks, Annotatability is aided by the trainability-aware graph embedding algorithm, which integrates information on annotation-trainability and gene expression to interpret cellular landscapes through the lens of any annotation of interest. Finally, by ranking cells according to their congruence with a given annotation, Annotatability identifies genes that are positively or negatively associated with an annotation of interest, e.g. a disease state.

Annotatability currently treats input annotations as nominal categories. For the special but common binary case (e.g. epithelial vs. mesenchymal, healthy vs. diseased tissue), the confidence in a cell’s annotation naturally gives rise to ordinal or numerical interpretations, where lack of confidence (and low variability) in one annotation may be considered as increased confidence in its complement. Moreover, Annotatability provides built-in tools for identifying intermediates between categories with ambiguous training dynamics scores, and for refining their discretization and ordering via trainability-aware graphs. In future work we aim to extend Annotatability to leverage the internal, potentially continuous, structure of the annotation set, if such exists, which in turn, will allow us to generalize both the re-annotation and graph embedding modules to incorporate the annotation structure. While the measures of confidence and variability of DNN predictions along the training process provided ample information to reveal cellular heterogeneity, structure, and other patterns related to label-incongruence, future work can incorporate additional measures derived from the training dynamics of a DNN to characterize additional aspects of the congruence of cells with their input annotations and potentially provide a wider interpretation lens into single-cell heterogeneity.

Here, we exemplified, through diverse applications and systems, the potential that annotation-trainability analysis holds for dissection, refinement, and interpretation of single-cell data. While we demonstrated the application of this type of analysis to single-cell RNA sequencing and spatial transcriptomics datasets, our approach is general, and in principle, it can readily extend to other single cell multi-omic techniques such as ATAC-seq[63] or Patch-seq[64], or in fact, to any biological domain where annotations are crucial for understanding complex biological phenomena, including but not limited to functional annotations of genes, taxonomic annotations of metagenomic sequences, pathway annotations in signaling networks, and interaction annotations in metabolic networks.

## 4 Methods

### 4.1 Annotatability workflow

Given input data which includes single-cell or spatial omics observables (e.g. scRNA-seq gene expression profiles) and corresponding annotations (e.g. cell type annotation per cell), Annotatability general-case workflow is as follows:

1. Preprocess the data.
2. Train a deep neural network (DNN) on the input data, monitoring the training dynamics and recording the prediction of the DNN after each epoch.
3. Calculate the training dynamics scores - confidence and variability.
4. classify each observable (e.g. cell) into one of three categories: easy-to-learn (correctly annotated), hard-to-learn (erroneously annotated) or ambiguous.
5. Optional step: filter out erroneously annotated cells.
6. Optional step: re-annotate erroneously annotated cells using the re-annotation module.
7. Optional step: construct trainability-aware graph embedding, integrating information from gene expression and training dynamics statistics, followed by signal-specific analysis.
8. Optional step: score and rank genes according to their (positive or negative) annotation trainability association.

#### Rationale

One of the main aims of Annotatability is to detect cells that are erroneously annotated and cells that are in ambiguous cell states. For both of those tasks, we train a deep-learning-based classifier to assign each cell to its input annotation and monitor the training dynamics. We set the model’s level of complexity to be sufficiently high to ensure that the network will eventually assign the input annotations to all provided examples; the underlying assumption is that a complex network such as the one we are using is capable of memorizing any input annotation, even if it is incorrect[65]. Formally, our approach is based on a model that involves selecting parameters to minimize empirical risk by stochastic gradient-based optimization procedure over *E* epochs. The model is assumed to establish a probability distribution over labels for a given observation (cell). The training dynamics of a given instance *i*, following previous work in the context of natural language processing[28], are described by statistical metrics computed over *E* epochs, which are then used as coordinates on a map. The first metric aims to quantify the level of confidence with which the learning algorithm assigns the correct label (*y*_*i*_) to an observation (the gene expression of the cell, *x*_*i*_) based on its probability distribution. Specifically, we define confidence, 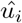, as the average probability assigned by the model to the correct label (*y*_*i*_) across all epochs:

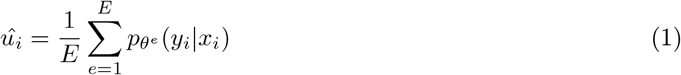

where 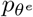 denotes the model’s probability with parameters *θ*^*e*^ for the e^*th*^ epoch. In addition, variability is defined using the standard deviation of 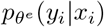 across epochs:

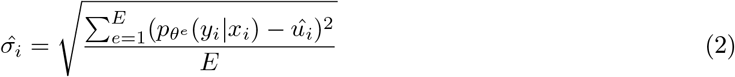

Based on previous studies in the fields of computer vision[66, 67] and natural language processing[28], our underlying assumption is that cells with high confidence and low variability (easy-to-learn) likely have correctly labeled annotations, which the network learns early in the training process. Conversely, cells with low confidence and low variability (hard-to-learn) are likely mislabeled and would be learned only at later stages of the training procedure. Cells with moderate confidence and variability are in an ambiguous/intermediate state associated with at least two different annotations.

### 4.2 Initial training phase

For training, we constructed a simple DNN consisting of 3 fully connected layers with ReLU activation[68]. As a loss function, we used cross-entropy loss, which is minimized using stochastic gradient descent (SGD) with an SGD optimizer. We used PyTorch[69] to perform the training procedure described above. To mitigate the effects of class imbalance (e.g. rare cell types) we used a weighted sampler, ensuring that rare annotations are given higher importance during the training process. The number of epochs is chosen to be sufficiently large, such that the empirical risk will be minimized and will reach a value close to zero by the end of the training procedure. The full procedure can be found in the publicly available code at: https://github.com/nitzanlab/Annotatability

### 4.3 Annotation-trainability score

We define the annotation-trainability positive association score (*g*_*j*_ per gene *j*), which utilizes training dynamics to identify genes with higher expression levels in cell states associated with a given annotation. Using the annotation-trainability association score, genes can be ranked based on the correlation between their expression in each cell and the corresponding confidence scores:

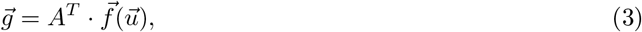

Where 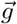 is the vector of gene scores, A is the gene expression matrix (rows correspond to cells and columns correspond to genes), *f* : *R* → *R* is a monotonically increasing function for rescaling confidence scores (by default, Annotatability uses the identity function *f*(*u*_*i*_) = *u*_*i*_), and 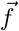 is a vector-valued function applying *f*(*x*) to every entry in the vector of cell confidence scores 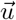. As a preprocessing step, the expression level of each gene is scaled to have variance 1 (L2 normalization).

Similarly, we define a annotation-trainability negative association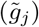, which identifies genes with lower expression levels in cell states associated with a given annotation based on training dynamics:

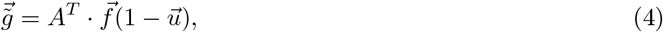

For the special case where 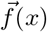 is the identity function, 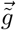 equals 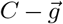 for some constant *C*, and thus, 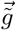 ranks genes in the reverse order relative to *g*. This order reversal holds for every non-zero linear function. However, for the general case where *f*(*x*) may be a non-linear monotonically increasing function, genes ranked by 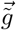 may be sorted differently from the reverse of their rankings by *g*.

### 4.4 Classifying cells based on confidence and variability scores

To classify cells as easy-to-learn, hard-to-learn, or ambiguous, Annotatability applies a threshold on the computed confidence and variability scores. This threshold is dataset-dependent, similarly to the case of area under the margin ranking statistic in the context of computer vision[26].

In the case that cells can be easily clustered in the Confidence-Variability plane to three groups corresponding to low-confidence/low-variability, mid-confidence/high-variability, and high-confidence/low-variability, a threshold can be set manually to distinguish the groups to hard-to-learn, ambiguous, and easy-to-learn, respectively. In the general case, however, we infer the corresponding confidence/variability thresholds using the following algorithm: (1) Randomly sample *c* cells (5% − 10%) and change their annotations (sample a different annotation out of the set of input annotations uniformly at random). (2) Next, train a DNN classifier (as described in Methods 4.2) and record the training dynamics. Set the threshold over the confidence and variability scores for hard-to-learn cells as the *q* percentile of the confidence and variability scores over the *c* reannotated cells. *q* is chosen to be between 25% and 90%, in a dataset-dependent matter.

The threshold between easy-to-learn cells and ambiguous cells is set, across all datasets, to Confidence score = 0.95, Variability score = 0.15. The threshold can also be tuned, keeping in mind that with more epochs the Confidence will increase, and the Variability will decrease.

### 4.5 Re-annotation of erroneously annotated cells

To correct the annotations of cells that were identified by Annotatability as erroneously annotated, a DNN classifier (as described in Methods 4.2) is trained exclusively on the subset of cells that were identified as correctly annotated. The erroneously annotated cells are then re-annotated according to the predictions of the newly trained DNN.

### 4.6 Trainability-aware graph embedding

To construct a trainability-aware gene expression graph, we first compute pairwise distances between cells by integrating gene expression information and training dynamics statistics. Specifically, the trainability-aware gene expression distance matrix 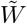 is computed as:

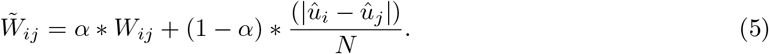

where *W* is the Euclidean distance matrix; *W*_*ij*_ is the Euclidean distance between cells *i* and *j* in gene expression space following dimensionality reduction using PCA (default number of principal components = 50), 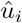 is the confidence score for cell *i, N* is the mean value of 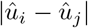 across all pairs 0 ≥*i, j* ≥ *n* (*n* is the total number of cells), and *α* is a tunable parameter 0 ≤*α* ≤1, which interpolates between a gene expression-based distance matrix (*α*=1) and a trainability-based distance matrix (*α*=0).

Next, the trainability-aware expression distance matrix 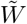 is transformed to an affinity matrix *M* using a Gaussian kernel:

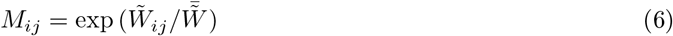

where 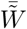 is the mean over all values in 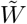.

Finally, the trainability-aware expression graph is constructed by computing a KNN graph (K = 15) over the affinity matrix *M*.

### 4.7 Data preprocessing

We used standard scRNA-seq preprocessing, which includes per-cell-normalization (to 10,000 counts), and a log transformation which is applied to stabilize variance (log(Normalized Expression Value + 1). For the EMT dataset, we use the raw datra. Therefore, we filtered out low-quality cells[10] by filtering out cells with high amounts of mitochondrial genes ( ≥ 5%) and cells with a high total count number (number of expressed genes higher than 4,000). In addition, we retained only highly variable genes (3,000).

## 5 Code availability

The code is publicly available at: https://github.com/nitzanlab/Annotatability

## 6 Data availability

The EMT and mSTZ scRNA-seq datasets were acquired from the Gene Expression Omnibus (GEO) database with the following accession numbers: EMT: GSE114687[49], mSTZ:GSE114687 [51] The PBMC data[70, 71] was downloaded using scvi-tools[32]. The MERFISH[39], 4i[42], seqFISH[43] and Visium[41] datasets were downloaded using Squidpy[72].

## A Supplementary Material

### A.1 Benchmarking

To benchmark the identification of erroneous annotations, we compared the results of Annotatability with the results of a state-of-the-art method: scReClassify[40], using two different classifiers: SVM and random forest. In addition, we compare the results to a KNN classifier, implemented in the same way as scReClassify. scReClassify: we implemented the method described in[40] in Python, and compared the inferred confidence vector by Annotatability to the inferred probability of the cell being misclassified by scReClassify, which is the cumulative probability of the cell belonging to cell types other than the corresponding input annotation.

For the comparison, we created a semi-synthetic setting, where we randomly perturbed the annotation of varying fractions of cells (10%, 20%, 30%, 40%, and 50%). We randomly subsampled each of the spatial datasets to 10,000 cells, with the exception of the Visium dataset which originally contained 2800 cells.

**Table 1:**
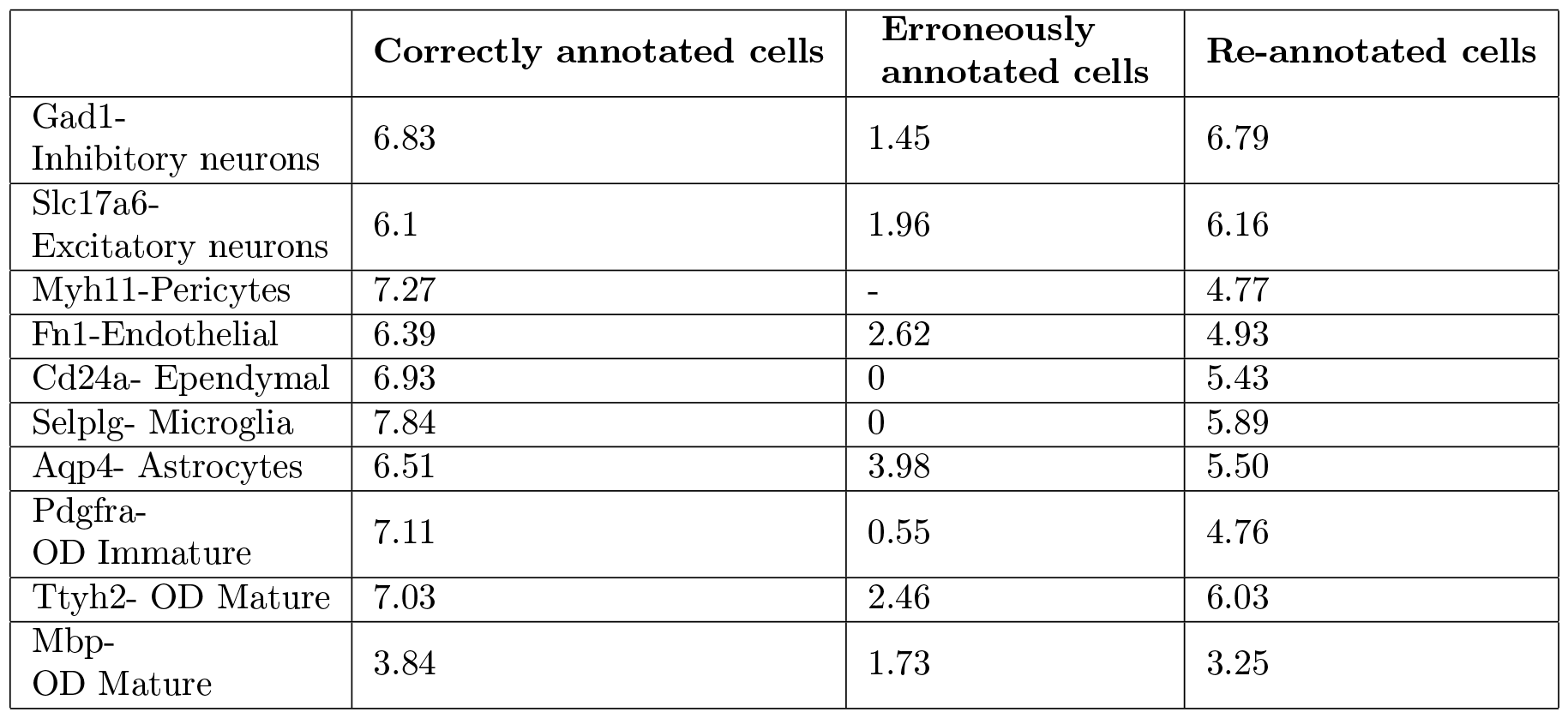
Mean expression of marker genes in correctly annotated cells, erroneously annotated cells and re-annotated cells in a MERFISH dataset of mouse hypothalamic preoptic region[39].

#### A.2 Supplementary Figures

**Supplementary Figure 1:**
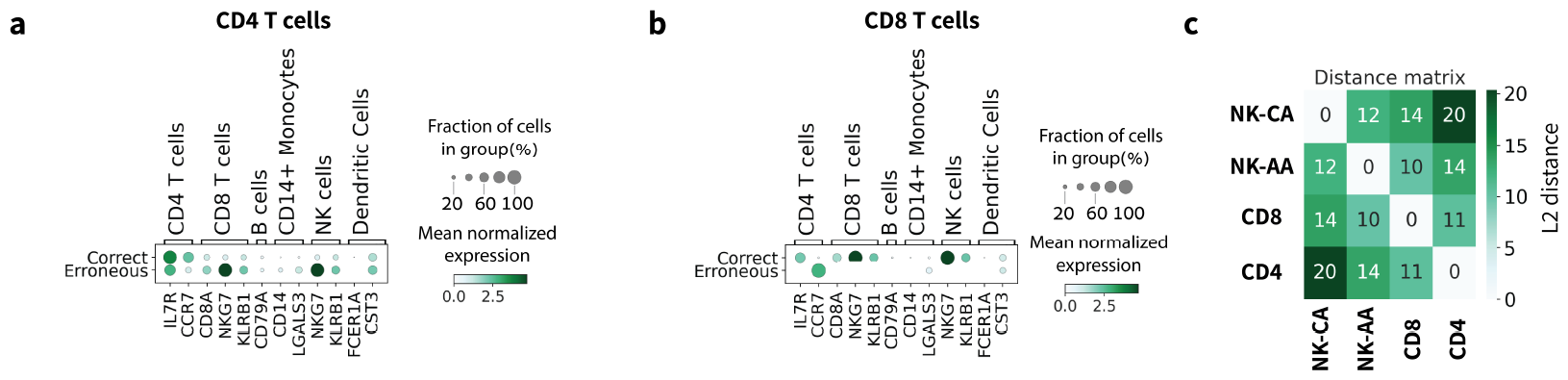
(a, b) Dot plots of the expression of cell type marker genes of the annotated cell types in either CD4 T cells (a) or CD8 T cells (b) identified as correctly annotated (top row) or erroneously annotated (bottom row) in human PBMCs scRNA-seq data. (c) L2 distance between the mean gene expression of the following four groups: NK cells which were classified as correctly annotated (CA), NK cells which were classified as ambiguously annotated (AA), CD8 T cells, and CD4 T cells.

**Supplementary Figure 2:**
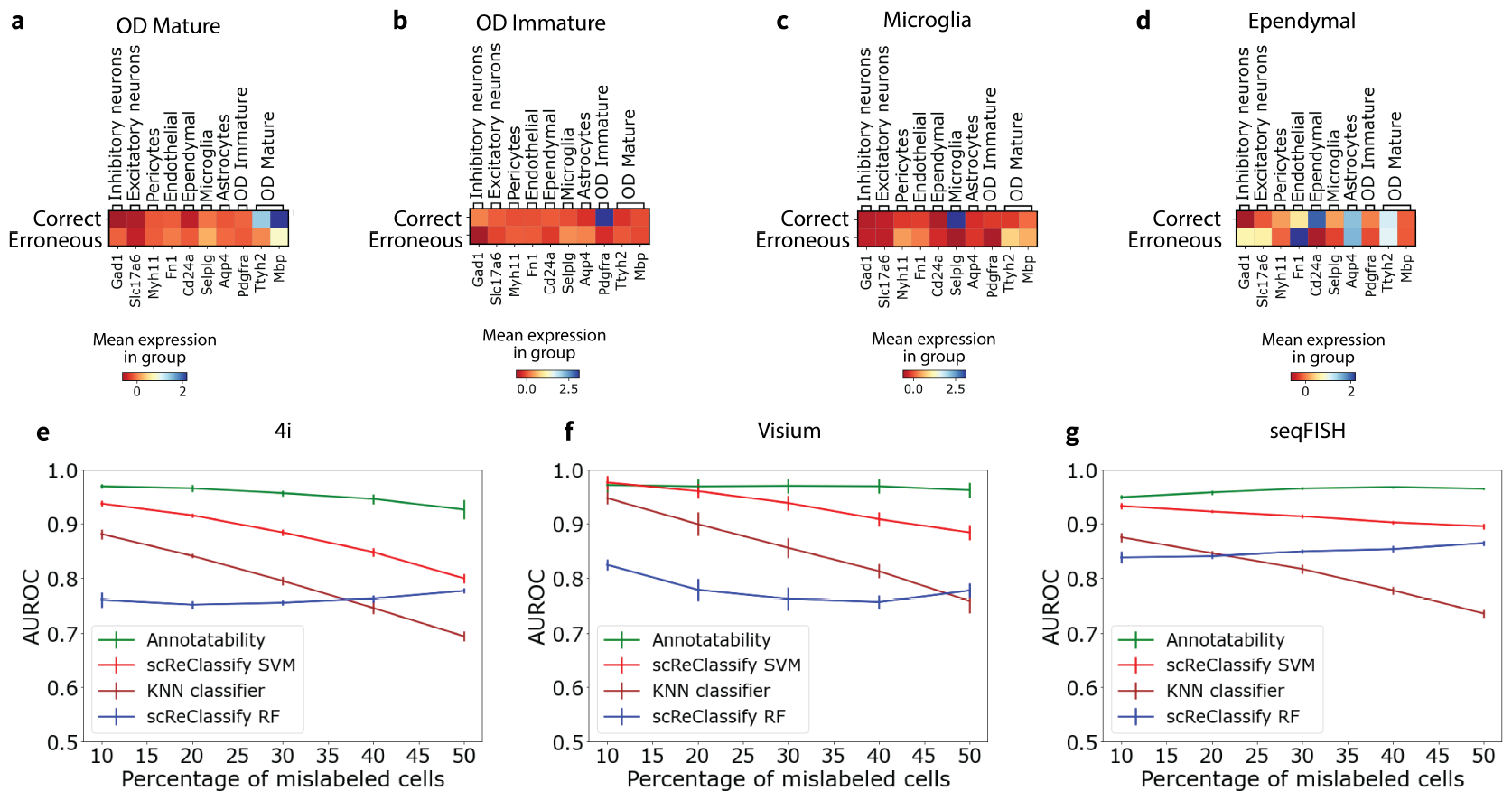
(a-d) Heatmaps of the mean expression of marker genes of the annotated cell types, in OD Mature (a), OD Immature (b), Microglia (c), and Ependymal cells (d) identified as correctly annotated (top row) or erroneously annotated (bottom row) in MERFISH dataset of mouse hypothalamic preoptic region[39]. The cell type marker genes[39]: *Gad1* (Inhibitory neurons), *Slc17a6* (Excitatory neurons), *Myh11* (Pericytes), *Fn1* (Endothelial), *Cd24a*(Ependymal), *Selplg* (Microglia), *Aqp4* (Astrocytes), *Pdgfra* (OD Immature), *Ttyh2, MBbp* (OD Mature). (e) AUCROC vs percentage of mislabeled cells, for three spatial datasets: Four i (43 genes), Visium (16562 genes), and Seqfish (351 genes), for four different methods: KNN classifier (brown), scReClassify with random forest classifier (blue), scReClassify with SVM classifier (red), and Annotatability(green).

**Supplementary Figure 3:**
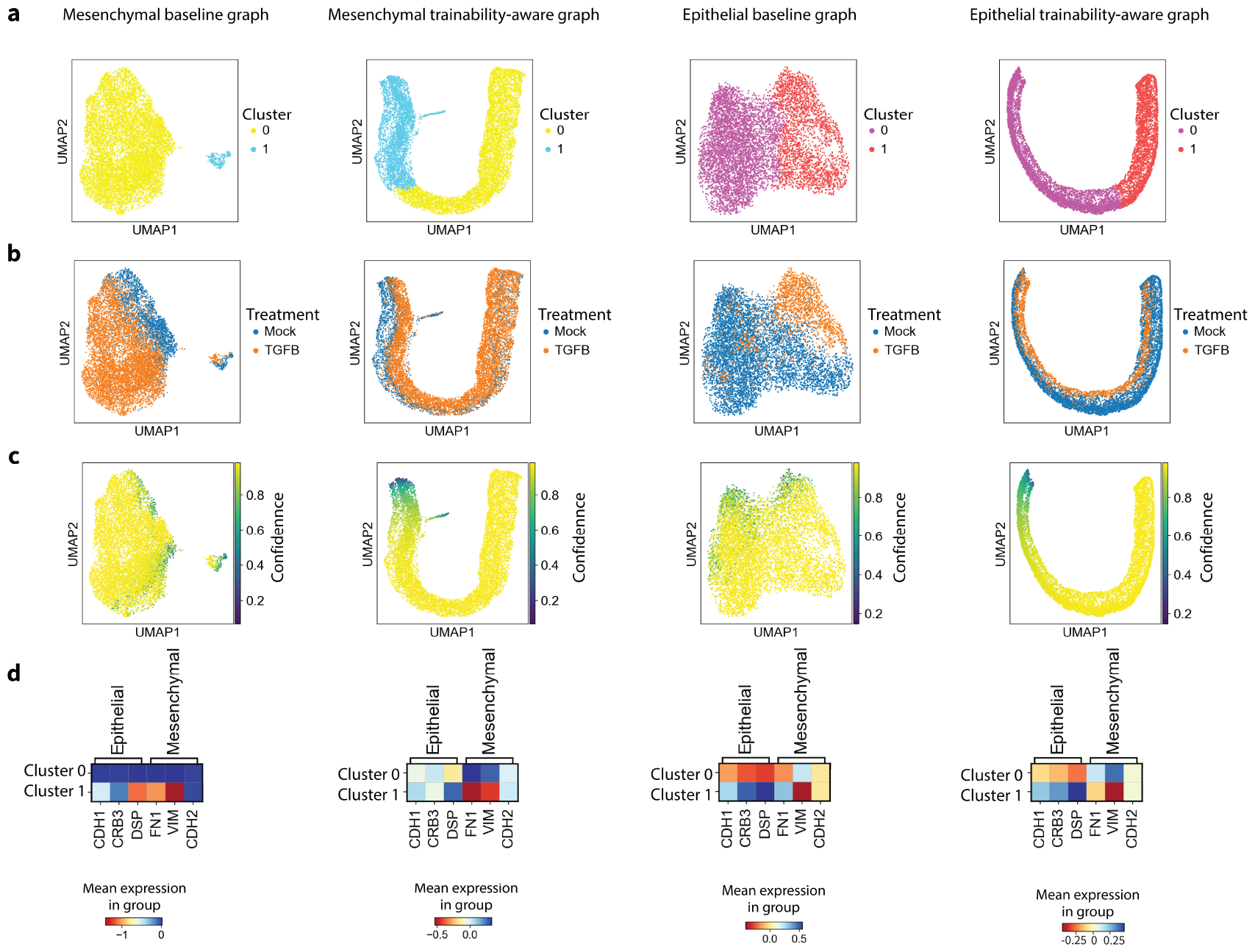
Analogous results to Figure 4g-j for HuMEC cell line in an EMT scRNA-seq data[49]. (a-d) 2D UMAPs of (left to right columns): baseline expression graph of Mesenchymal annotated cells, trainability-aware graph of Mesenchymal annotated cells, baseline expression graph of Epithelial annotated cells, and trainability-aware graph of Epithelial annotated cells. The graphs are colored by (top to bottom rows): Louvain clustering, treatment, inferred confidence by Annotatability. (d) Heatmaps of scaled and centered expression of Epithelial and Mesenchymal marker genes in cells belonging to the two clusters as in (a). The columns, from left to right, correspond to the graphs as in (a-c).

**Supplementary Figure 4:**
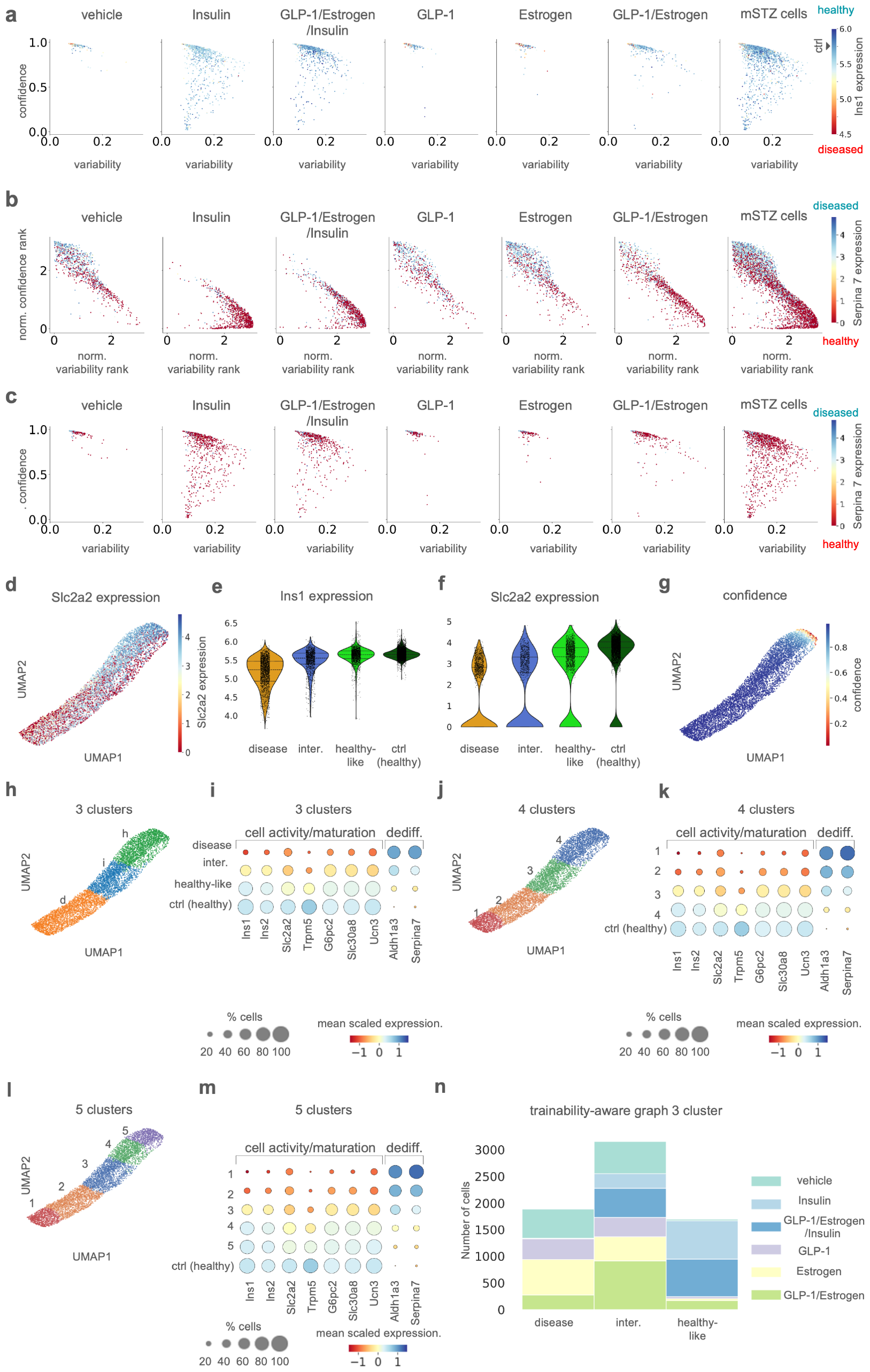
a) 2D map of the confidence and variability scores with respect to a disease annotation colored by *Ins1* expression, for cells from mSTZ diabetic models receiving different follow-up treatments (six panels starting from the left), and across all mSTZ cells regardless of follow-up treatment (rightmost panel). Supplementary Figure 4: b) 2D map of the normalized ranks of the confidence and variability scores with respect to a disease annotation colored by *Serpina7* expression, for the same sets of cells as in (a). c) Same as the 2D map in (a), colored by *Serpina7* expression. d) 2D UMAP showing the trainability-aware graph for all mSTZ cells, colored by *Slc2a2* expression. e) Violin plots showing the distribution of *Ins1* expression in cells from each of the three Louvain clusters obtained from the trainability-aware graph of the mSTZ cells, or from the healthy control. f) Violin plots as in (e) for *Slca2a2* expression. g) 2D UMAP as in (d), colored by confidence scores for all mSTZ cells. h) 2D UMAP as in (d), colored by three Louvain clusters. i) Dot plots showing the mean scaled expression of the same marker genes as in Figure 5c for the three Louvain clusters as in (h) vs. control healthy cells (bottom row) j) Same 2D UMAP as in (h), shown for four clusters; the number of clusters was increased by tuning the Louvain clustering hyperparameters. k) Same dot plots as in (i) for four Louvain clusters. l) Same 2D UMAP as in (h), shown for five clusters. m) Same dot plots as in (i) for five Louvain clusters. n) Stacked bar plot showing the number of cells in each of the three Louvain clusters for each treatment.

